# Chemically synthesized guide RNAs can direct CRISPR-CasRx cleavage of circRNAs with high efficiency and specificity

**DOI:** 10.1101/2022.08.30.505797

**Authors:** Karim Rahimi, Maria Schertz Andersen, Sabine Seeler, Thomas Birkballe Hansen, Jørgen Kjems

**Author notes:** The authors contributed equally to the manuscript (Karim Rahimi, Maria Schertz Andersen). Correspondence (J. Kjems); (K. Rahimi).

## Abstract

Circular RNAs (circRNAs) are characterized by a covalently closed circular structure, formed from pre-mRNAs through an alternative splicing mechanism named back-splicing. CircRNAs have been shown to play a regulatory role in the development of eukaryotic organisms and to be implicated in human diseases. However, the extensive sequence-overlap between circRNAs and their linear RNA counterparts makes it technically difficult to deplete circRNAs without affecting their linear host, which complicates functional studies. Therefore, it is important to identify the most efficient and specific strategy for circRNA depletion. In this study, we demonstrate that CRISPR/RfxCas13d (CasRx)-mediated circRNA depletion is, for the circRNAs studied, more efficient than Argonaute 2-dependent short hairpin RNA (agoshRNA)-mediated depletion and with fewer off-target effects on the linear host RNAs. Furthermore, we show that synthetic guide RNAs (syn-gRNAs) can be used in combination with CasRx to efficiently deplete circRNA, ciRS-7. Finally, none of the knockdown (KD) strategies tested (pre-gRNA, gRNA, syn-gRNA and agoshRNA) showed any significant off-target effects on the global transcriptome. Taken together, CasRx-mediated circRNA KD strategies, using either vector-based or syn-gRNA, are useful tools for future studies on circRNA functions.

## Background

CircRNAs are formed from pre-mRNAs via back-splicing events and have been reported to play significant functions in cell signalling and the development of eukaryotic organisms (1). They are expressed in a highly tissue-specific manner (2, 3) and known to be dysregulated in cancer and neurodegenerative disorders (Kristensen *et al*., 2018; Gokool *et al*., 2020; Kristensen *et al*., 2021). The most well characterized functions of circRNA include, but are not limited to, microRNA sponging (6–8), protein sponging (9, 10) and transcription regulation (11). Compared to linear RNA species, circRNAs are more stable presumably due to their resistance to exonucleases (2, 12, 13).

CircRNAs are known to be expressed under the control of the same regulatory elements as their host gene (14) and usually share the same exon sequences. Only the back-splicing junction (BSJ) is unique to the circRNA which makes it difficult to design loss-of-function studies that specifically target the circRNAs without affecting the co-expressed linear RNA. Different strategies such as small interfering RNAs (siRNAs) and short hairpin RNAs (shRNAs) have been used to deplete circRNAs; however, these methods may induce significant off-target effects, not only on the linear host RNA of the corresponding circRNA, but also on the transcriptome in general (15, 16).

The development of the Clustered Regularly Interspaced Short Palindromic Repeats (CRISPR) technology has made a remarkable revolution in biological studies. In the first CRISPR studies, DNA was targeted using the Cas9 enzyme guided by a complement guide RNA (gRNA) (17–19), but with the advent of the CRISPR-Cas13 system, directed cleavage of RNA targets also became possible (20–23). Konermann *et al*. introduced CasRx as one of the most efficient Cas13 enzymes to target RNA molecules (24). The CasRx KD setup is based on expressing the CasRx enzyme along with an expression cassette for either precursor guide RNA (pre-gRNA) or mature gRNA. The pre-gRNA primary transcript contains a spacer sequence, which is complementary to the targeted RNA molecule and flanked by two short direct repeat elements that generate stem-loop structures (Figure 1A). After transcription, the pre-gRNA is processed by CasRx to create the mature gRNA containing the 5’ direct repeat followed by a 14-26 nucleotide (nt) spacer sequence (24, 25).

**Figure 1:**
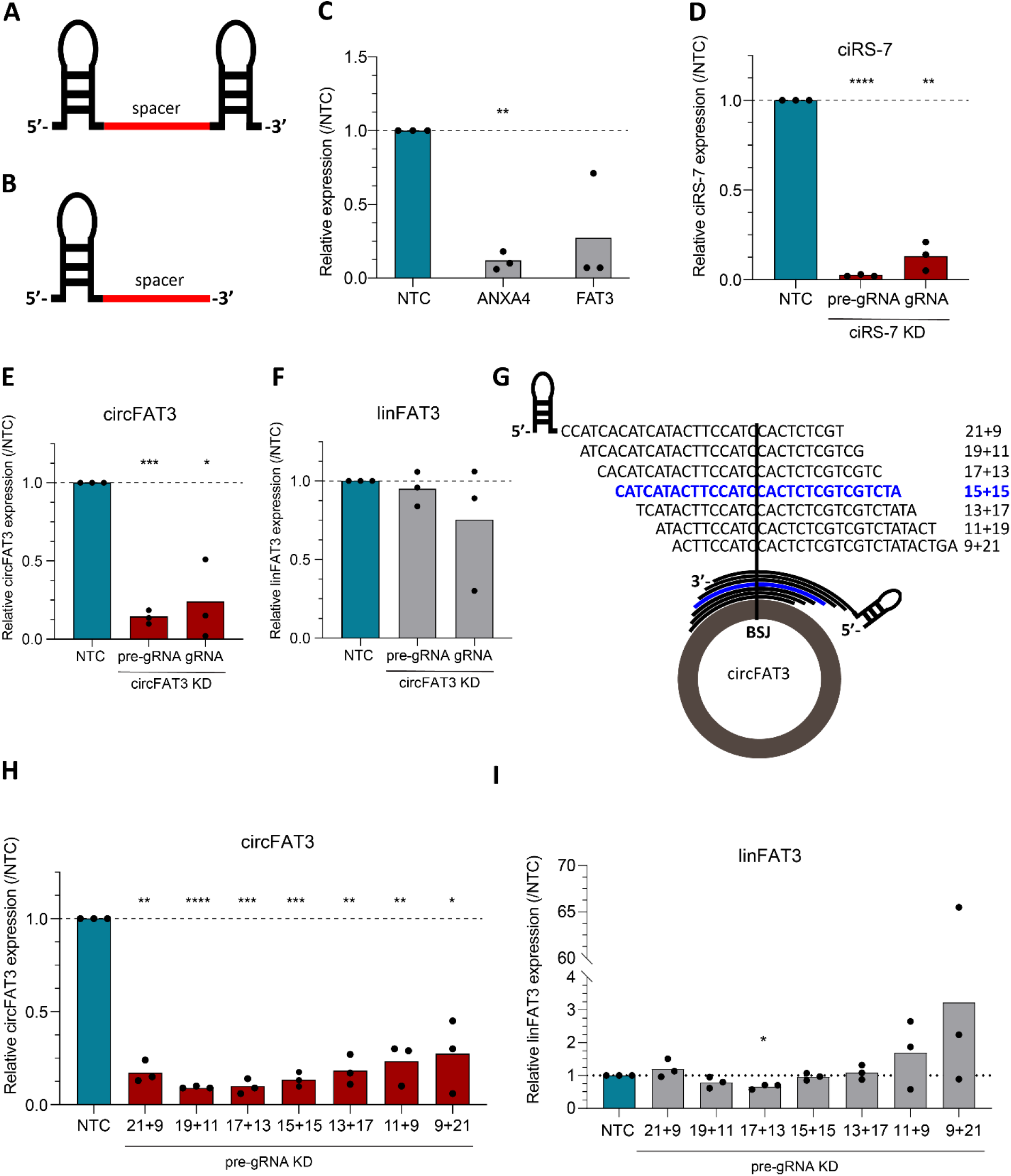
Optimization of vector-based CasRx circRNA knockdown strategies. A) The pre-gRNA primary transcript forms a scaffold containing a spacer flanked by two direct repeat elements. Upon binding to CasRx, the 3’ direct repeat is cleaved off, leaving the 5’ direct repeat and 14-26 nt spacer as the mature gRNA. B) The primary transcript of the fixed size gRNA expression cassette, only contains the 5’ direct repeat followed by the fixed size spacer corresponding to the mature gRNA. This transcripts will not be further processed. C) Quantification of *ANXA4* and *FAT3* mRNAs by RT-qPCR upon pre-gRNA-based KD. D) RT-qPCR showing pre-gRNA- and gRNA-mediated KD of circular RNA, ciRS-7. E-F) Quantification of circFAT3 (E) and linFAT3 (F) by RT-qPCR upon pre-gRNA- and gRNA-mediated KD. G) Design of seven different spacers of equal 30 nt length shifted across the BSJ of circFAT3. Numbers denote number of nucleotides complementary to up- and downstream sequences relative to the BSJ. H-I) RT-qPCR showing 21+9, 19+11, 17+13, 15+15, 13+17, 11+19 and 9+21 nt spacer mediated KD of circFAT3 (H) and the effect on linear *FAT3* (I). RT-qPCR expression was normalized to *GAPDH*; bars show mean fold change compared to NTC (set to 1); all experiments were performed in biological triplicates; light grey bars show linear RNA expression; white bars show circRNA expression; one sample t-test, *p<0.05, **p<0.01, ***p<0.001, ****p<0.0001; NTC: non-targeting control; KD: knockdown; BSJ: back-splicing junction.

Konermann *et al*. demonstrated that expression of the mature CasRx-gRNA from a plasmid gives rise to a fully functional gRNA containing the 5’ direct-repeat followed by a fixed size spacer (Figure 1B). The study also reported that guide RNA with spacers longer than 20 nt significantly targeted and depleted RNA molecules containing the complementary sequence. More recently published studies have demonstrated efficient CasRx-based KD of circRNA KD using both pre-gRNA and gRNA approaches (26, 27).

Over the last few years, synthetic guide RNAs for DNA targeting with CRISPR-Cas9 have been introduced (28). These type of gRNA molecules are chemically synthesized and do not require any cloning steps prior to transfection, making them easy to use in the laboratory. In addition, it was previously shown that chemically modified syn-gRNAs give rise to less inflammatory signaling and are thus less toxic for mammalian cells than vector-based CRISPR Cas9 approaches (29). Therefore, their application in biological studies have increased dramatically. A recent study showed that chemically modified syn-gRNAs in combination with CasRx could efficiently deplete linear RNAs in human cells (30). However, to the best of our knowledge, there is no published report using syn-gRNAs for targeting circRNAs.

Comparative studies have shown that the vector-based CasRx systems generally show a higher KD efficiency and fewer off-targets compared to shRNAs (24, 26, 27). However, a more specific Dicer-independent shRNA-processing pathway, which mimics the processing of miR-451 (31, 32), has been developed (for review, see (33)). By shortening the shRNA stem to 16-19 bp, Dicer cleavage is prevented, which leads to cleavage by Argonaute-2 (AGO2) between nucleotides 10-11 of the 3’-passenger strand (34). The sliced RNA intermediate is further processed by poly(A)-specific ribonuclease (PARN), producing the mature guide strand RNA (35). This pathway prevents entry of the passenger strand into the RNA-induced silencing complex (RISC) and reduces the associated off-target effects. Because of the AGO2-dependent processing mechanism, the shRNAs were named agoshRNAs or Dicer-independent shRNAs (dishRNAs) (34, 36, 37). AgoshRNAs have previously been used to target a single specific circRNA to assess its functional impact during neural development (38), but no direct comparison with the Cas13 system has been conducted.

In this study we compared the circRNA-specific KD efficiency of agoshRNA and CasRx pre-gRNA and investigated off-target effects on the linear RNA. Furthermore, we investigated KD efficiency using synthetic pre-gRNA and gRNA molecules targeting the BSJ of the circRNA ciRS-7, also known as CDR1as (39). Finally, we used total transcriptome profiling to compare the global off-target effects of agoshRNA-, pre-gRNA-, gRNA-as well as syn-gRNA in connection with specific circRNA KD.

## Results

### Optimization of vector-based CasRx-mediated circRNA knockdown strategies

As an initial proof of concept for the CasRx-based KD, we tested a vector-based pre-gRNA design against the linear *ANXA4* and *FAT3* mRNAs (Supplementary table 1). Human neuroblastoma cells (SH-SY5Y) were co-transfected with the pcDNA3_CasRx-GFP vector and the respective pre-gRNA expression vector and total RNA was harvested 48 hours (hrs) post-transfection. For all CasRx-based KD experiments in our study, we included a previously described non-targeting control (NTC) sequence without known RNA targets in the human transcriptome (Supplementary Table 1) (24). Using quantitative reverse transcription PCR (RT-qPCR), we found that both pre-gRNAs significantly deplete *ANXA4* and *FAT3* with an average efficiency of 88.6±3.4% (mean±SEM) and 71.6±21.1%, respectively (Figure 1C).

To compare the KD efficiency of pre-gRNA and gRNA, their respective expression vectors were designed to target the BSJ of ciRS-7 (circBase ID: hsa_circ_0001946) and circFAT3 (circBase ID: hsa_circ_0000348) utilizing a 30 nt spacer encompassing 15 nt of the flanking regions on either side of the BSJ (Supplementary Tables 1-2). We predicted that this design would only target BSJ-containing RNA because pre-gRNAs carrying a spacer of 15 nt or shorter have previously proven unable to degrade the targeted RNA molecule, even if fully complementary (24). To test this hypothesis, SH-SY5Y cells were co-transfected with one plasmid expressing the CasRx fused to GFP (pcDNA3_CasRx-GFP vector) and another expressing pre-gRNA or gRNA (Addgene; #109053 and #109054). Based on RT-qPCR with divergent primers amplifying the BSJ, the KD efficiency for ciRS-7 and circFAT3 amounted to 97.6±0.3% and 86.7±2.6% using the pre-gRNA and 86.7±4.8% and 77.1±14.5% using the gRNA, respectively (Figure 1D-E). For the gRNA, we observed an off-target KD of linear *FAT3* (24±22.1%), which was only 3.5±6.6% for the pre-gRNA (Figure 1F). Based on these results, we decided to proceed with the pre-gRNA setup.

To investigate the importance of target position relative to the BSJ, the KD of circRNA and off-target effects on the corresponding linear RNA were measured for vector-based pre-gRNAs with 30 nt spacers targeting seven different regions, all encompassing the circFAT3 BSJ (21+9, 19+11, 17+13, 15+15, 13+17, 11+19 and 9+21 nt; Numbers indicate the length of the 5’ and 3’ target sequence relative to the BSJ; Figure 1G) in conjunction with CasRx expressing vector. Co-transfection of the CasRx and pre-gRNA expression vectors resulted in a significant KD for all pre-gRNA variants (Figure 1H; one-sample t-test). Especially the 19+11, 17+13 and 15+15 designs resulted in efficient circFAT3 KD (90.9±0.6%, 90.3±2.4% and 86.7±2.6%, respectively; Figure 1H). The 15+15 design showed little effect on the linear *FAT3* transcript (3.5±6.6% KD; Figure 1I), while the 19+11 and 17+13 designs showed a stronger off-target effect on linear *FAT3* (KD of 21.4±9.6% and 33.8±4.5%, respectively (Figure 1I)). We therefore decided to continue with the 15+15 nt design to target additional circRNAs and to compare the KD efficiency side-by-side with agoshRNAs.

### Vector-based CasRx pre-gRNA-mediated circRNA knockdown is more efficient than agoshRNA-mediated knockdown

Previous studies have investigated the optimal agoshRNA design for efficient transcription and AGO2-processing (40, 41). Based on this, we decided to use agoshRNAs with a stem length of 17 nt, a loop of 4 nt and an A-C mismatch below the stem (Figure 2A). To directly compare the KD efficiency and off-target effect of vector-based pre-gRNAs and agoshRNAs on circRNAs and the corresponding linear transcripts, we designed pre-gRNAs and agoshRNAs targeting the BSJ of five selected circRNAs ranging from high to low expression in SH-SY5Y cells including ciRS-7 (circBase ID: hsa_circ_0001946), circFAT1 (circBase ID: hsa_circ_0001461), circFAT3 (circBase ID: hsa_circ_0000348), circNRIP (circBase ID: hsa_circ_0004771) and circTET1 (circBase ID: hsa_circ_0093996); (Figure 2B). The pre-gRNA spacers were based on the 15+15 nt design (pre-gRNA-NTC included as a control) and the agoshRNAs were likewise designed to target the circRNAs in the middle of the BSJ with a 10+11 nt design (Figure 2A); a 21 nt version of the NTC was included as a control. We determined the expression level of the circRNA, using divergent primers amplifying the BSJ, and their respective linear counterpart using convergent primers spanning the adjacent exon-exon junction (Figure 2B; grey bars), except for ciRS-7, which shows negligible amounts of linear product (6, 39, 42). All RT-qPCR products were validated by Sanger sequencing.

**Figure 2:**
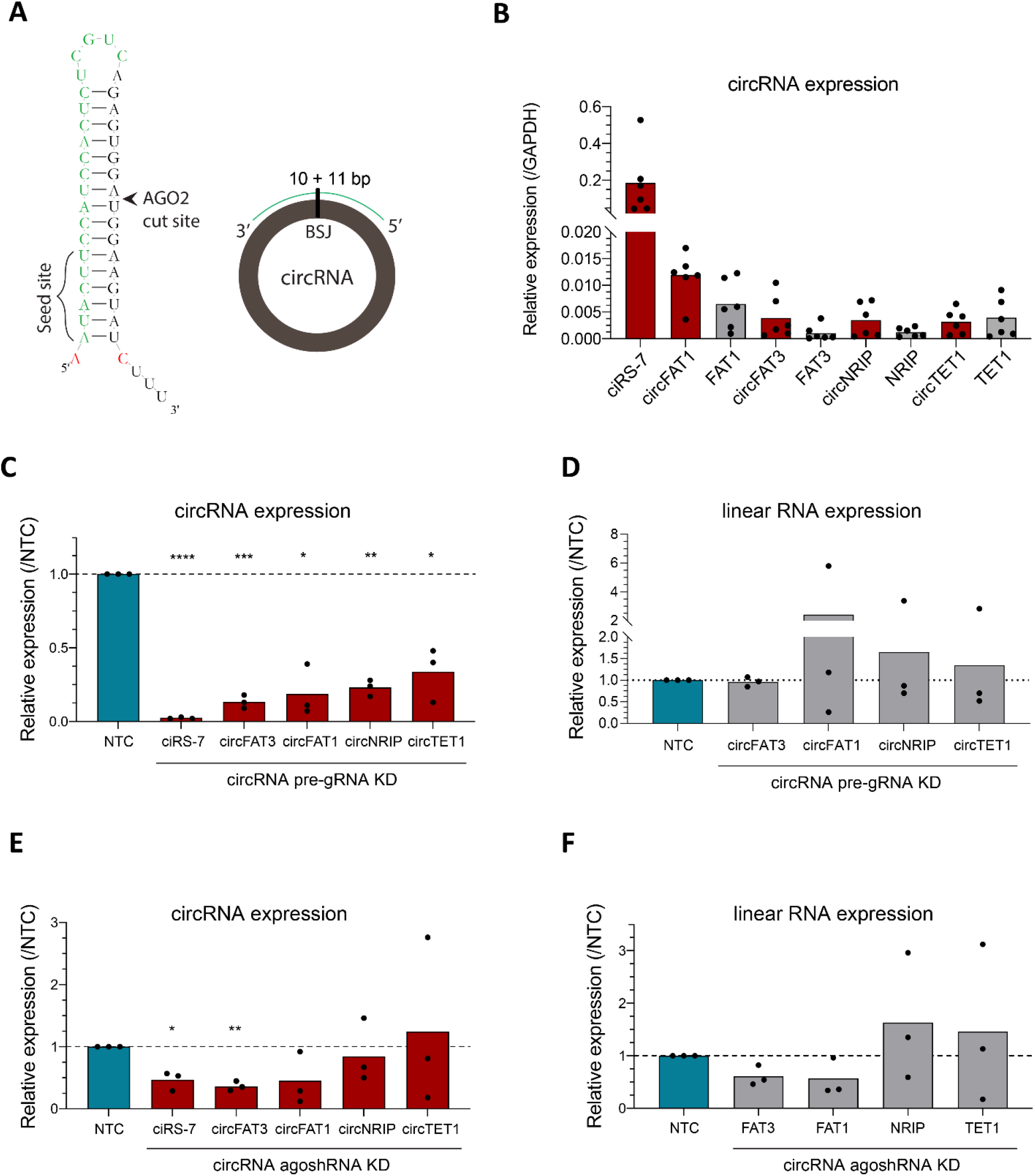
Comparing CasRx/pre-gRNA and agoshRNA-mediated circRNA knockdown. A) Predicted agoshRNA structure (left side); green nucleotides indicate the mature guide strand RNA, which can target the BSJ of the circRNAs, as indicated to the right; red nucleotides indicate A-C mismatch below the stem; the seed site is marked with brackets. B) Expression of selected circRNAs and their linear counterpart in SH-SY5Y cells determined by RT-qPCR; expression was normalized to *GAPDH*. C-D) CircRNA KD efficiency (C) and effect on linear RNA (D) mediated by pre-gRNA. E-F) CircRNA KD efficiency (E) and effect on linear RNA (F) mediated by agoshRNA. Bars show mean fold change (relative expression) compared to NTC (set to 1); all experiments were performed in biological triplicates; light grey bars show linear RNA expression; white bars show circRNA expression; one sample t-test, *p<0.05, **p<0.01, ***p<0.001, ****p<0.0001; NTC: non-targeting control; KD: knockdown; BSJ: back-splicing junction.

The use of CasRx-directed pre-gRNAs led to a more efficient KD of all circRNAs tested, ranging from 97.6±0.3% KD for ciRS-7 to 66.2±10.7% KD for circTET1 (Figure 2C), compared to agoshRNA-mediated circRNA KD, which ranged from 64.7±4.1% KD for circFAT3 to 24.8±77.6% upregulation for circTET1 (Figure 2E). The pre-gRNA-mediated approach resulted in relative low levels of linear RNA off-targeting, specifically ranging from 3.5±6.6% KD for *FAT3* to 140.9±171.1% increase for *FAT1* (Figure 2D), while the agoshRNA had a more pronounced effect on the linear host RNAs, ranging from 44.7±20.5% KD for *FAT1* to 63.3±70% upregulation for *NRIP* (Figure 2F). However, despite the increased variation seen for agoshRNA-dependent KD none of the effects on linear RNA expression were statistically significant (Figure 2F). We conclude, CasRx-pre-gRNAs generally showed a better KD efficiency and lower off-target effect on linear host RNAs compared to the agoshRNAs.

### Synthetic pre-gRNAs and gRNAs lead to efficient knockdown of ciRS-7

To investigate whether the CasRx-mediated KD of circRNAs is compatible with the use of synthetic guide RNA molecules, we chemically synthesised gRNA (syn-gRNA) (Figure 3A) and pre-gRNA (syn-pre-gRNA; Figure 3B) with a ciRS-7-specific 15+15 nt design. As a negative control, a synthetic non-targeting control guide RNA (syn-gRNA-NTC) was designed. Furthermore, all the syn-gRNAs were stabilised by including 2’-O-methyl (Me) analogs and 3’ phosphorothioate (S) internucleotide linkages in the first and last three RNA residues (Figure 3A-B). To investigate if the chemical modifications affected the KD efficiency, an unmodified syn-gRNA targeting the ciRS-7 BSJ (syn-gRNA_non-mod) was included in the experimental setup (Supplementary Table 3). To investigate the effect of spacer length in the syn-pre-gRNA setup, spacers of 30 nt (15+15; syn-pre-gRNA) and 22 nt (11+11; syn-pre-gRNA_short) were designed to target the BSJ of ciRS-7 (Figure 3B). SH-SY5Y cells were first transfected with the pcDNA3_CasRx-GFP vector and six hours later with the respective syn-gRNA. Total RNA was harvested at 16, 24 and 48 hrs post-transfection and quantified with RT-qPCR as described above.

**Figure 3:**
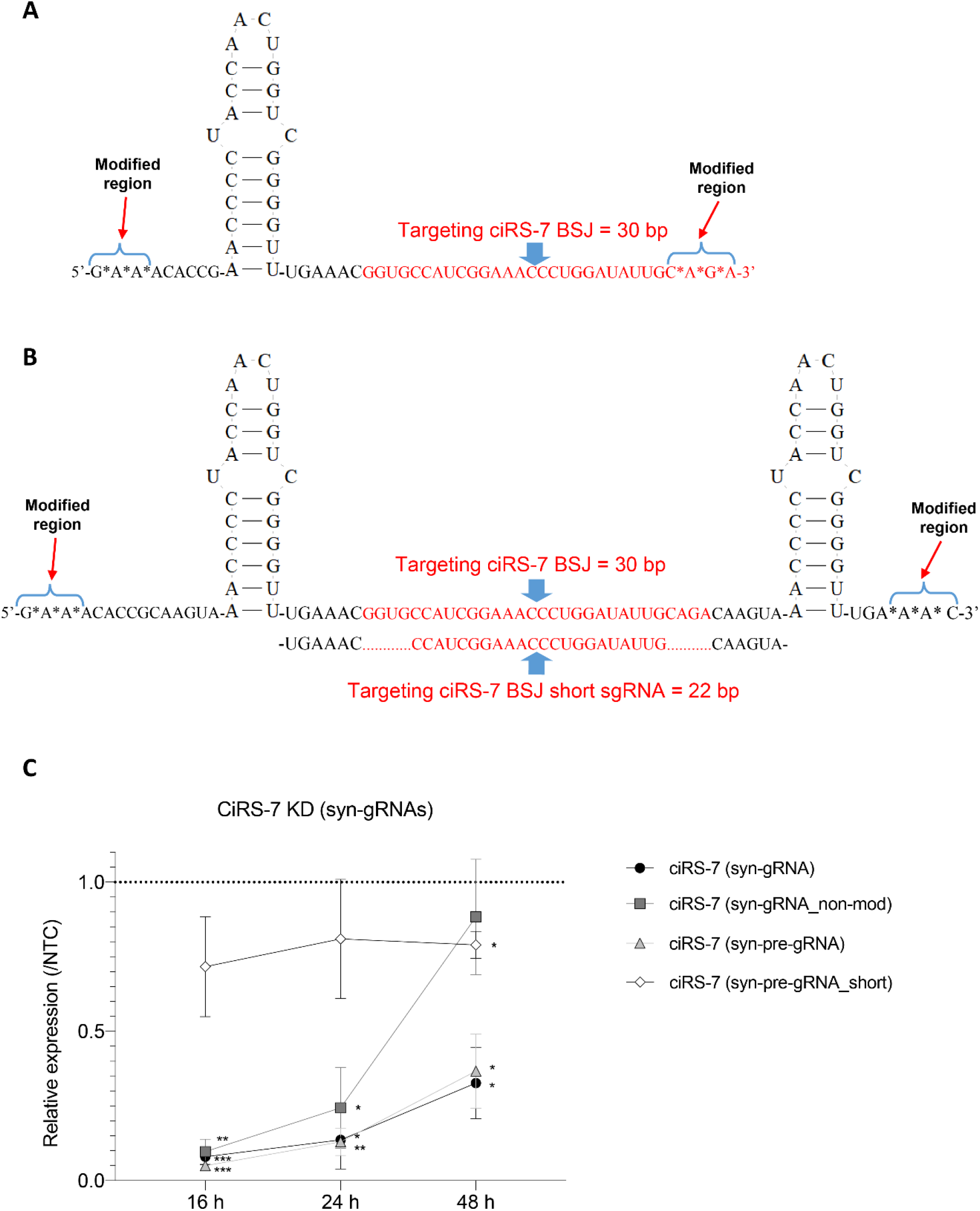
Synthetic gRNA and pre-gRNAs targeting the BSJ of ciRS-7 lead to efficient knockdown. A) The syn-gRNA contains one direct repeat element followed by a 30 nt spacer targeting the ciRS-7 BSJ. For syn-gRNA-NTC, the ciRS-7 spacer was replaced with 30 nt NTC. The first and last three nucleotides of the RNA marked with an asterisk are replaced with 2’-O-methyl analogs with 3’ phosphorothioate internucleotide linkages. B) The syn-pre-gRNA design contains two direct repeats flanking the ciRS-7 BSJ targeting spacer; the RNA modifications are the same as in panel A. For the syn-pre-gRNA setup, ciRS-7 targeting spacers of either 22 or 30 nt were designed. The former design is referred to as syn-pre-gRNA_short. C) CiRS-7 depletion at 16, 24 and 48 hrs post-transfection using syn-gRNAs and syn-pre-gRNAs and with and without modification. Expression levels of ciRS-7 were determined by RT-qPCR with divergent primers targeting the BSJ. Bars show mean fold change ±SEM compared to NTC set to 1; all experiments were performed in biological triplicates; bars show circRNA expression; one sample t-test, *p<0.05, **p<0.01, ***p<0.001; NTC: non-targeting control; KD: knockdown; BSJ: back-splicing junction.

At 16 hrs post syn-gRNA transfection, ciRS-7 expression was reduced by 92±2.65% and 90±4.18% using either syn-gRNA or syn-gRNA_non-mod in comparison to syn-gRNA-NTC, respectively (Figure 3C). This was also observed for the syn-pre-gRNA with a 30 nt spacer (95±1.53% KD; Figure 3C). The syn-pre-gRNA with a 22 nt spacer only showed a 28.3±16.7% KD at this time point which further decreased to 21±4.5% KD efficiency 48 hrs after transfection (syn-pre-gRNA_s; Figure 3C). While syn-gRNA_non-mod resulted in efficient KD of 90±4.18% and 75.7±13.5% at 16 and 24 hrs post-transfection, this decreased to 11.7±19.4% KD 48 hrs post transfection. However, the modified versions of both syn-gRNA and syn-pre-gRNA with 30 nt spacer size resulted in 67.3±11.9% and 63.3±12.5% KD even 48 hrs post-transfection (Figure 3C).

We conclude that the chemically modified syn-gRNA showed better KD at later timepoints compared to the non-modified version. Likewise, the long syn-pre-gRNA (30 nt spacer) resulted in a more pronounced KD than the short version (22 nt spacer; syn-pre-gRNA_short). The syn-gRNA and syn-pre-gRNA showed similar KD efficiencies at all timepoints.

### Neither CasRx-nor agoshRNA-based KDs give rise to pronounced off-target effects when targeting ciRS-7

To test if the KD strategies used here could lead to more general off-target effects, we subjected samples from each ciRS-7 KD (pre-gRNA, gRNA, syn-gRNA and agoshRNA) and the corresponding NTC controls to mRNA sequencing, generating 30 to 35 million reads per sample. Differential expression analysis was performed for each KD setup versus NTC using DESeq2. However, we observed no significant differentially expressed genes (DEGs) in any of the experimental setups when using the adjusted p-value (p.adj <0.05) (Supplemental Figure 1A-D), indicating that none of the setups tested trigger significant off-target effects at the transcriptomic level.

Finally, with no significant off-targets for any of our four KD approaches, we went on to investigate possible regulatory effects of ciRS-7 KD on mRNA expression in SH-SY5Y cells. We hypothesized that overlapping genes between the DEG lists from the four KD approaches would give us high confidence biological targets upon ciRS-7 KD. Comparing the overlap between DEG lists with non-adjusted p <0.01, we found no differentially expressed and overlapping transcripts that were shared between at least two of the KD experiments, indicating that ciRS-7 KD does not have any major effect on the transcriptome in undifferentiated SH-SY5Y cells after 48 hrs of depletion.

## Discussion

Loss-of-function studies are essential to determine the function of a gene of interest. The extensive sequence overlap between circular and linear RNAs makes it technically difficult to deplete only the circRNA form without affecting their linear host RNA. Therefore, it is important to identify the most efficient and specific strategy for circRNA depletion. Here, we demonstrated that vector-based CasRx-mediated circRNA depletion was, for the circRNAs studied, more efficient than agoshRNA-mediated depletion and with less pronounced effects on the linear host RNAs. The high efficiency and specificity of the CasRx-mediated vector-based KD system aligns well with two recently published papers on CasRx-based KD of circRNAs in comparison to shRNAs (26, 27).

In our experimental setup, we found no evidence for significant off-targets across the transcriptome neither via CasRx- nor agoshRNA-mediated ciRS-7 depletion. Notably, Zhang *et al*. found 25 off-target genes for an shRNA targeting a circularized form of *EGFP*, compared to no off-target genes using CasRx-mediated KD (27). Furthermore, the first published CasRx paper for linear RNAs found 542 and 915 off-target genes upon shRNA KD of *B4GLANT1* and *ANXA4*, respectively, but no significant ones for the CasRx-mediated KD of the same genes (24). The apparent higher specificity for agoshRNA compared to shRNA is in accordance with previous comparisons of these methods and may be explained by the more selective incorporation of the 5’ target strand (33, 43). However, since the target sequences used differ between each KD study, they may also have a different potential for causing off-targets.

Our data show for the first time that synthetic syn-gRNAs can be used in combination with CasRx to target a BSJ of circRNA, here ciRS-7. We found that the modified versions of both syn-gRNA and syn-pre-gRNA more effectively depleted ciRS-7 at later time points (up to the 48 hrs tested), presumably due to enhanced stability in the cytoplasm. Neither the vector- nor the syn-gRNA-mediated ciRS-7 KD strategy showed significant off-target effects at the transcriptomic level. The use of syn-gRNAs for circRNA depletion has several advantages: they do not require cloning steps, can be synthesized at the same cost as siRNAs and display no apparent off-target effects. We found that syn-gRNA and syn-pre-gRNA were equally efficient in depleting the circRNA of interest and we therefore recommend to use syn-gRNA given the shorter final length and lower synthesis costs.

Furthermore, our syn-gRNA study showed that the 30 nt syn-pre-gRNA was more efficient than the shorter form (22 nt), also in line with the results published by Zhang, *et al*., where the authors found the vector-based gRNA with spacers from 24-30 nt to be most efficient (27). In contrast, a recently published study used syn-gRNAs in combination with CasRx and a 23 nt spacer sequence and obtained efficient KD of multiple linear target genes (30). Méndez-Manilla *et al*. also investigated linear RNA KD efficiency upon chemical modification of the syn-gRNAs, showing that Me-, S- and Me-S-modified gRNAs were more efficient than the non-modified forms (30), in line with our findings at later time points. Additionally, they showed that the modified syn-gRNAs showed extensive KD at 24, 48 and 96 hrs after nucleofection, in contrast to unmodified syn-gRNAs which lost their function after 48 hrs (30). This aligns well with our study, where the KD of ciRS-7 for the non-modified syn-gRNA declined between 24 and 48 hrs.

## Conclusions

Taken together, the two previously published studies (26, 27) as well as our study on CasRx-mediated circRNA KD show that the CasRx setup generally appears more efficient than shRNAs or agoshRNAs with no or little off-target effects on linear host genes, which makes it superior for future circRNA KD studies. In terms of off-target effects on other genes, the agoshRNA setup seems to match the CasRx-based protocol. Additionally, and for the first time, we used and validated optimally designed circRNA-specific chemically synthesized syn-gRNAs for significant circRNA depletion, even 48 hrs post-transfection. These modified syn-gRNAs were able to significantly deplete a highly expressed circRNA (ciRS-7) in SH-SY5Y cells without any off-target effect at the transcriptomic level. This potentially results in replacing the conventional methods of circRNA functional analysis approaches including siRNA- and shRNA-based technologies. This is because syn-gRNA setup has higher sensitivity in particular for circRNA-specific assays, and it is a feasible technique that costs the same or less than those aforementioned strategies which make it a very interesting technology for the RNA community. Future studies will show if this technique is able to target circRNAs in vivo with novel therapeutic avenues.

## Methods

### Plasmid constructs

#### RfxCas13d (pcDNA3_CasRx-GFP) construct

The RfxCas13d-GFP cassette was amplified from the plasmid pXR001:EF1a-CasRx-2A-EGFP (Addgene #109049) using the primers: 5’-AAGAATGGTACCCGTACGGCCACCATGAGC (FW) and 5’-AAGAATGGGCCCAGGTCTCGAGGTCGACGGTA (RE). The “Phusion Hot Start II DNA Polymerase” (Thermo Fisher, MA, USA) was used for PCR amplification. The obtained PCR product was digested with KpnI and ApaI, gel purified and ligated into the linearized pcDNA3 vector using the aforementioned restriction enzymes. This new CasRx-GFP cassette was driven by the CMV promoter and contained the WPRE translation enhancer element. The pcDNA3_CasRx-GFP vector was Sanger sequenced and co-transfected with either pre-gRNA or gRNA expression vectors for KD experiments.

#### Pre-gRNA / gRNA design and constructs

Both pre-gRNA (Addgene #109054) and gRNA (Addgene #109053) expression vectors were purchased from Addgene. The pre-gRNA vector contains two direct repeat elements downstream of the U6 promoter. There are two BbsI digestion sites between the repeat elements which results in two sticky ends. The gRNA expression vector only contains one direct repeat element followed by two BbsI restriction sites. Both pre-gRNA- and gRNA-based spacers were designed as two separate oligos (Supplementary tables 1-2) and synthesized by Sigma Aldrich. The oligos were annealed, which resulted in sticky ends compatible with the linearized pre-gRNA and gRNA vectors using the BbsI restriction enzyme.

#### AgoshRNA design and constructs

The pKHH030 vector (Addgene #89358) contains two BbsI restriction sites downstream of the U6 promoter and is typically used for gRNA expression. Here, the pKHH030 vector was modified for agoshRNA expression. For this purpose, the transcriptional start site was moved one base downstream and changed from G to A, which is more efficient for AGO2-processing (34). In addition, the nucleotide position after the agoshRNA insertion site was modified to a C to ensure an A-C mismatch at the bottom of the agoshRNA stem. Finally, a transcriptional termination signal (TTTTTTT) was inserted immediately after the C. In short, part of the region immediately downstream of the U6-promoter was PCR amplified from the pKHH030 vector with the following primers: 5’-GGGACAGCAGAGATCCAGTT-3’ (FW) and 5’-TAATGGATCCGAAAAAAAGAGGTCTTCTCGAAGACACGGTGTTTCGTCCTTTCCACA-3’ (RE) and NotI/BamHI digested. The digested product was ligated back into the NotI/BamHI linearized vector. For agoshRNA constructs, single stranded DNA oligos with BbsI overhangs were annealed and inserted into the BbsI-linearized empty vector (oligos are listed in Supplementary table 4).

### Synthetic guide RNAs design

Syn-pre-gRNA and syn-gRNA were designed to mimic the expressed pre-gRNA and gRNA from the corresponding plasmid constructs targeting the BSJ of ciRS-7. A synthetic non-targeted control guide RNA (syn-gRNA-NTC) was designed to be used as negative control. For the syn-pre-gRNA design, two different spacer sizes were designed with either 30 or 22 nts length. To protect the syn-pre-gRNA and syn-gRNA molecules from exonuclease degradation, three nucleotide bridges at both 5’ and 3’ ends of the RNA oligos were modified by 2’-O-methyl analogs and 3’ phosphorothioate internucleotide linkages. For the syn-gRNA design, an extra oligo batch was synthesized using the same sequence except that it remained non-modified (syn-gRNA_non-mod). All syn-gRNA oligos were synthesized and quality controlled by Synthego (SYNTHEGO, CA, USA). All syn-gRNA oligos are listed in Supplementary table 3.

### Cell maintenance

SH-SY5Y cells were maintained in Dulbecco’s modified Eagle’s media (DMEM/F12) (Invitrogen, GIBCO, MA, USA) supplemented with 10% fetal bovine serum (FBS; Invitrogen, GIBCO, MA, USA) and 1% Penicillin-Streptomycin (P/S) (Thermo Fisher Scientific, MA, USA). The cells were passaged before reaching 90% confluency using 0.05% trypsin (Thermo Fisher Scientific). Cells were cultured at 37°C and 5% CO2.

### Transfections

#### CasRx, pre-gRNA, gRNA and agoshRNA expressing plasmids

SH-SY5Y cells were transfected at a confluency of 70-80%. For CasRx experiments, the pcDNA3_CasRx-GFP vector (2 µg per 6-well) was co-transfected with either pre-gRNA or gRNA expression plasmids (1 µg per 6-well) using Lipofectamine 3000 kit (5 µl Lipo 3000 and 5 µl P3000 reagents per 6-well; Thermo Fisher Scientific) according to the manufacturer’s guidelines. AgoshRNA expression vectors (2.5 µg per 6-well) were transfected using Lipofectamine 3000 kit (7.5 µl Lipo 3000 and 5 µl P3000 reagents per 6-well; Thermo Fisher Scientific) according to the manufacturer’s instruction. After overnight incubation with the transfection reagents, a media change with fresh DMEM/F12 media (10%FBS, 1%P/S) was performed. RNA samples were harvested 48 hours post-transfection.

#### CasRx and synthetic gRNAs

Syn-gRNAs were diluted according to manufacturer’s instructions. Briefly, pellets were resuspended in 1X TE Buffer (Synthego, CA, USA) to a concentration of 100 µM for storage at -20°C. Directly before usage, the syn-gRNA solution was further diluted 1:10 using nuclease-free water. One day prior to transfection, SH-SY5Y cells were seeded with 8*10^5^ cells/cm^2^. The following day, the cells were first transfected with the pcDNA3_CasRx-GFP plasmid (2.5 µg per 6-well) using Lipofectamine 3000 kit (5 µl Lipo 3000 and 5 µl P3000 reagents per 6-well; Thermo Fisher Scientific) according to the manufacturer’s guidelines. After six hours of incubation and shortly before starting the second round of transfection, the media containing the initial transfection reagents was changed to fresh maintenance media. The syn-gRNA oligos and syn-gRNA-NTC were transfected using Lipofectamine RNAiMAX transfection reagent (5 µl per 6-well; Thermo Fisher Scientific) following the manufacturer’s instructions. Different concentrations ranging from 15-175 nM for the syn-gRNAs were initially tested (data not shown). All main experiments were performed using a final concentration of 58 nM for all the five syn-gRNAs separately. Twelve hours after the second transfection, the media containing the transfection reagents was replaced with fresh DMEM/F12 media (10%FBS, 1%P/S). RNA was collected at 16, 24, and 48 hours after the second transfection round. All transfection were performed in biological triplicates.

### RNA extraction

RNA extraction was started by lysing the cells in TRIzol reagent (Invitrogen) and followed by the usage of Direct-zol RNA Miniprep Kit (ZYMO RESEARCH, CA, USA) according to the manufacturer’s instructions including DNase on-column treatment for 15 min at room temperature. RNA concentrations and purity were determined using Nanodrop Lite (Thermo Fisher Scientific).

### cDNA synthesis and RT-qPCR

Complementary DNA (cDNA) for RT-qPCR gene expression analysis was synthesized from 1 µg of total RNA using the SuperScript VILO cDNA Synthesis Kit (Invitrogen) according to the manufacturer’s guidelines. Quantitative PCR was performed with Platinum SYBR Green qPCR SuperMix (Invitrogen) on LightCycler 480 (Roche) according to standard procedures, using divergent primer sequences spanning the back-splicing junction for circRNAs or convergent intron-spanning primers for linear RNAs (primer sequences are listed in Supplementary table 5). Expression levels were normalized to *GAPDH*. All RT-qPCR expression analyses were performed in 3 technical replicates.

### Library preparation and poly(A) RNA sequencing

RNA concentrations were measured using the Qubit RNA HS Assay Kit (Thermo Fisher Scientific) on a Qubit 4 Fluorometer (Thermo Fisher Scientific), as well as using Nanodrop Lite (Thermo Fisher Scientific). RNA quality was checked using Agilent Bioanalyzer 2100 (Agilent Technologies) and all RNA integrity (RIN) values exceeded 8.0. Library preparation, and poly(A) RNA sequencing was performed by Beijing Genomics Institute (BGI) in Copenhagen. Briefly, 1 µg total RNA input was used. Non-stranded and polyA-selected RNA library preparation was applied and PE100 sequencing was performed following the specific protocol for sequencing on a DNBSEQ sequencing platform (MGI, Shenzhen, China). Library size and purity control was performed on a Agilent 2100 Bioanalyzer (Agilent Technologies) using a High-Sensitivity DNA-Chip.

### Computational analysis of poly(A) RNA sequencing

Filtering and trimming of mRNA-sequencing reads was performed using trim_galore (version 0.6.5dev). The processed reads were mapped to the human reference genome (hg19) using STAR (version 2.7.3a)(44) with default settings. The mapped reads were then quantified per gene annotation using featureCounts (45) with gene annotation from Gencode (version 28). Differential expression analysis (DEA) was based on the featureCounts count data using DESeq2 (46) in R (http://www.r-project.org).

### Statistical analysis

The RT-qPCR dataset was analysed using GraphPad Prism version 9.0.2. One-sample t-tests assuming unequal variance with a theoretical mean of 1.0 for NTC samples were used for fold change comparisons.

### Data availability

The data is deposited to the Gene Expression Omnibus (GEO) repository database with the accession number GSE189963.

## Acknowledgements

We thank Anne Færch Nielsen for critical input and reading of the manuscript. Library preparation and poly(A) RNA sequencing was performed by Beijing Genomics Institute (BGI) in Copenhagen.

## Author contributions

The author contributions were written using the CrediT taxonomy (47). **K.R**.: conceptualization, methodology, formal analysis, investigation, writing – original draft, visualization, supervision, project administration; **M.S.A**.: conceptualization, methodology, formal analysis, investigation, writing – original draft, visualization; **S.S**.: conceptualization, methodology, formal analysis, investigation, writing – review and editing, visualization; **T.B.H**.: formal analysis, writing – review and editing; **J.K**.: conceptualization, resources, writing – review and editing, supervision, project administration, funding acquisition.

## Supplementary information

**Supplementary Figure 1:**
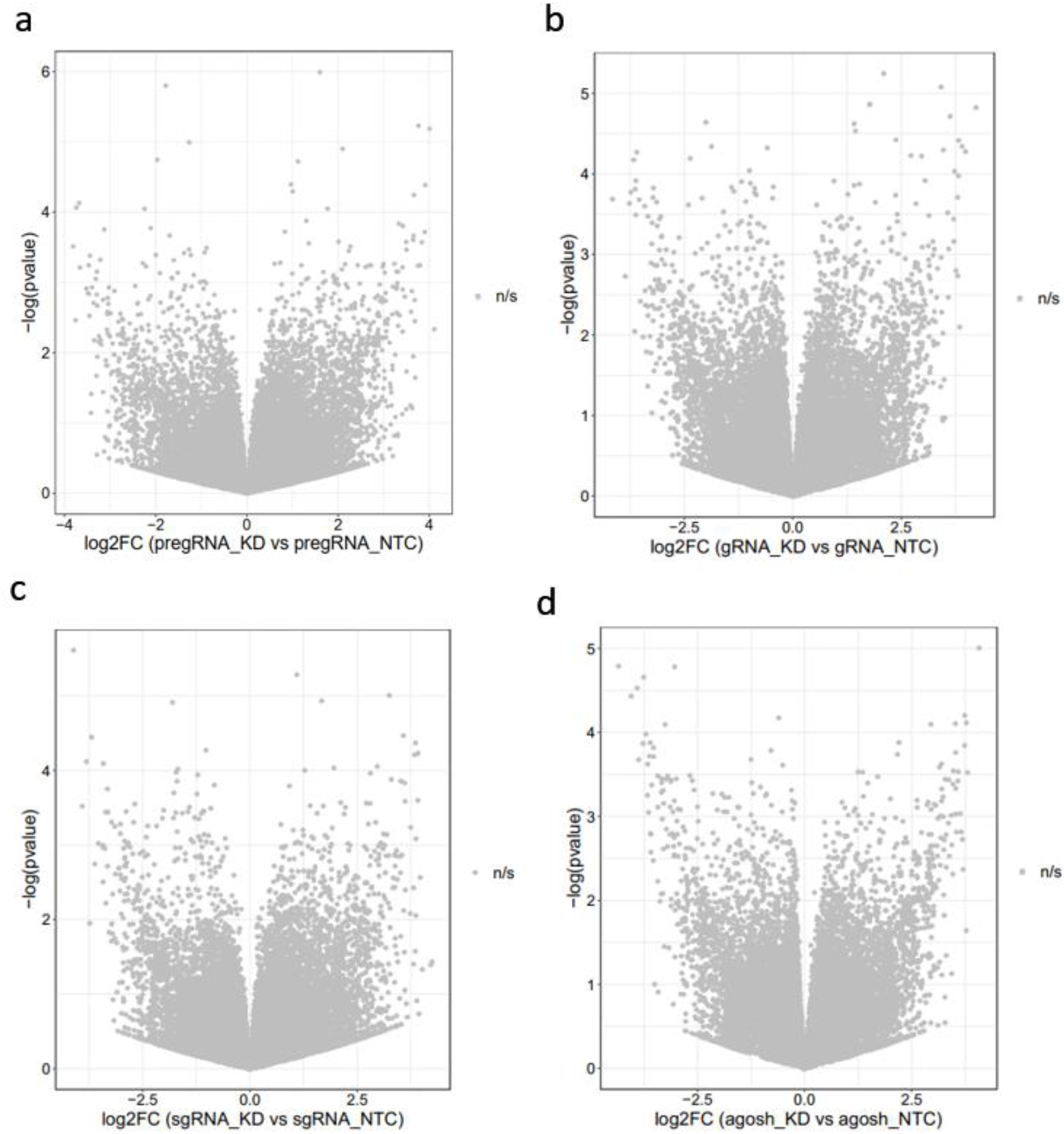
Neither CasRx- nor agoshRNA-based ciRS-7 knockdown strategies give rise to pronounced off-target effects. A-D) Volcano plots showing differential gene expression analysis (DEA) for ciRS-7 KD experiments using pre-gRNA (A), gRNA (B), syn-gRNA (C) and agoshRNA (D). RNA-seq was performed in three biological replicates for each ciRS-7 KD setup and for each matched NTC, respectively. DEA was performed using DESeq2; log2-fold change (log2FC) between NTC and KD are shown in the x-axis; -log10(p-values) are shown in the y-axis. KD: knockdown, DEGs: differentially expressed genes, NS: not significant.

**Supplementary table 1.**
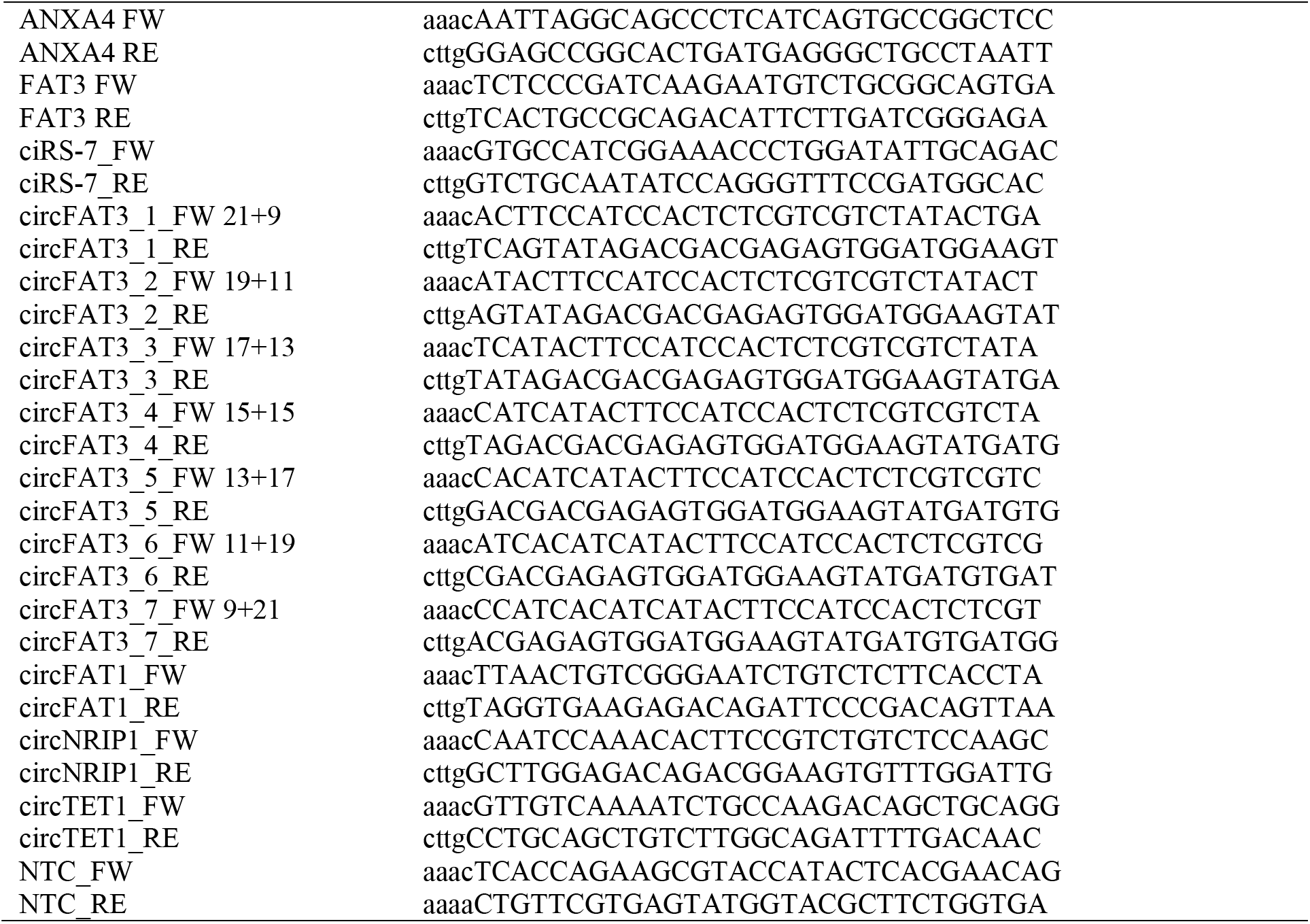
pre-gRNA oligos (5’>3’)

**Supplementary table 2.**
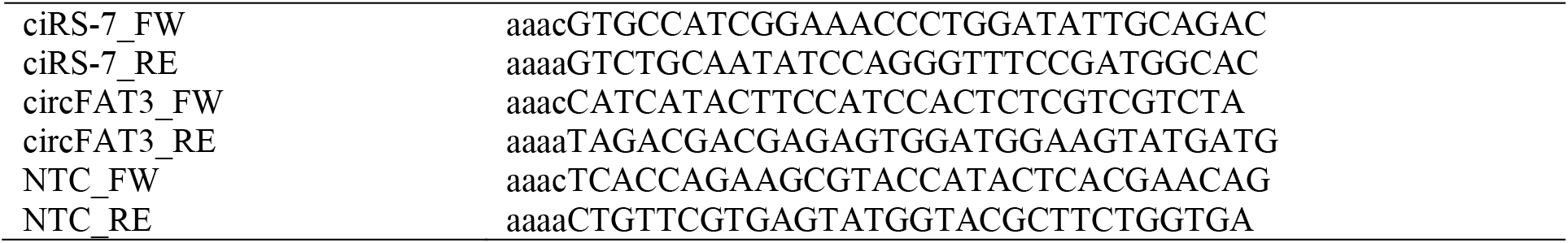
gRNA oligos (5’>3’)

**Supplementary table 3.**
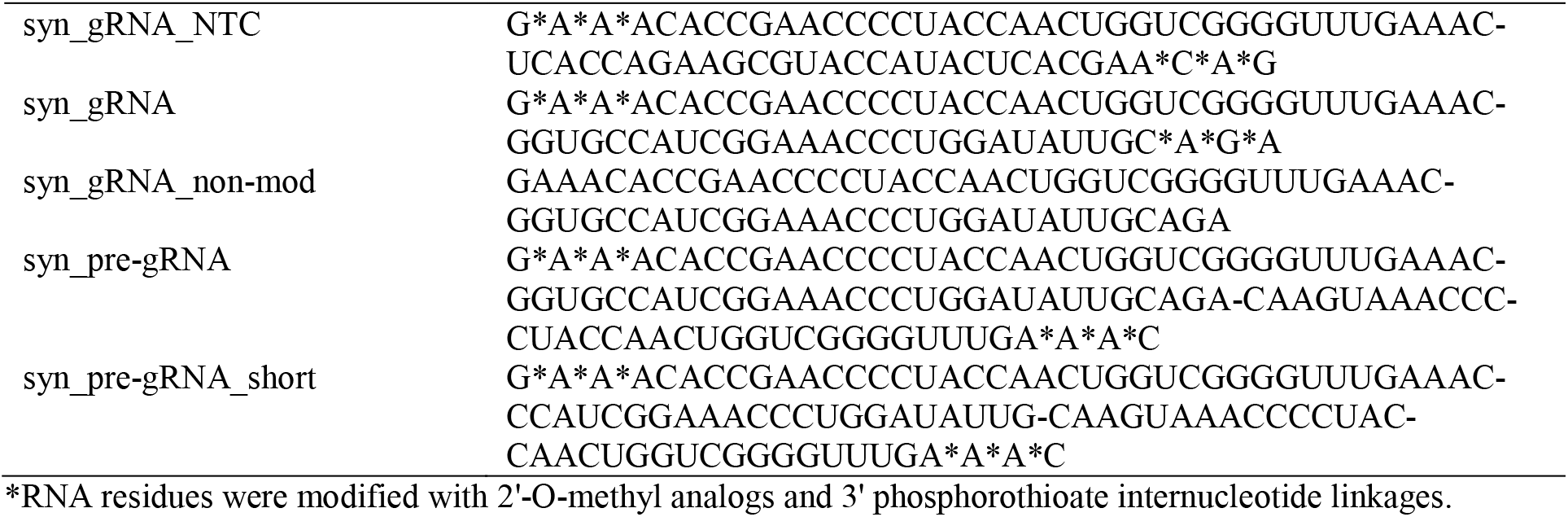
synthetic guide RNA oligos (syn-gRNAs) (5’>3’)

**Supplementary table 4.**
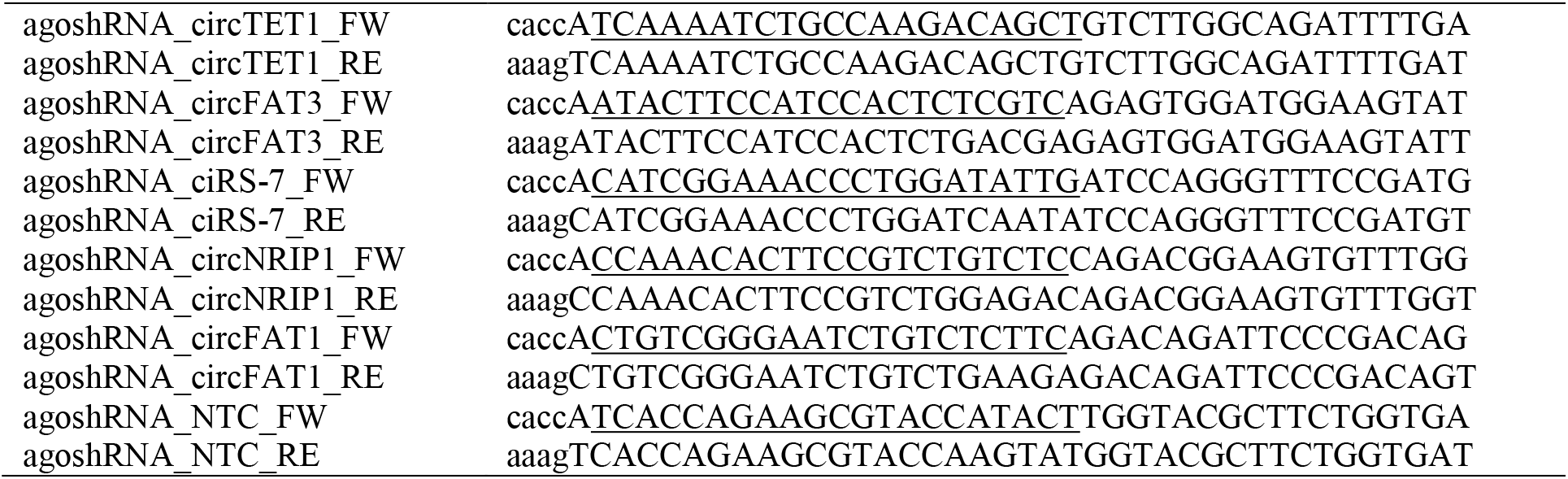
agoshRNA oligos (5’>3’)

**Supplementary table 5.**
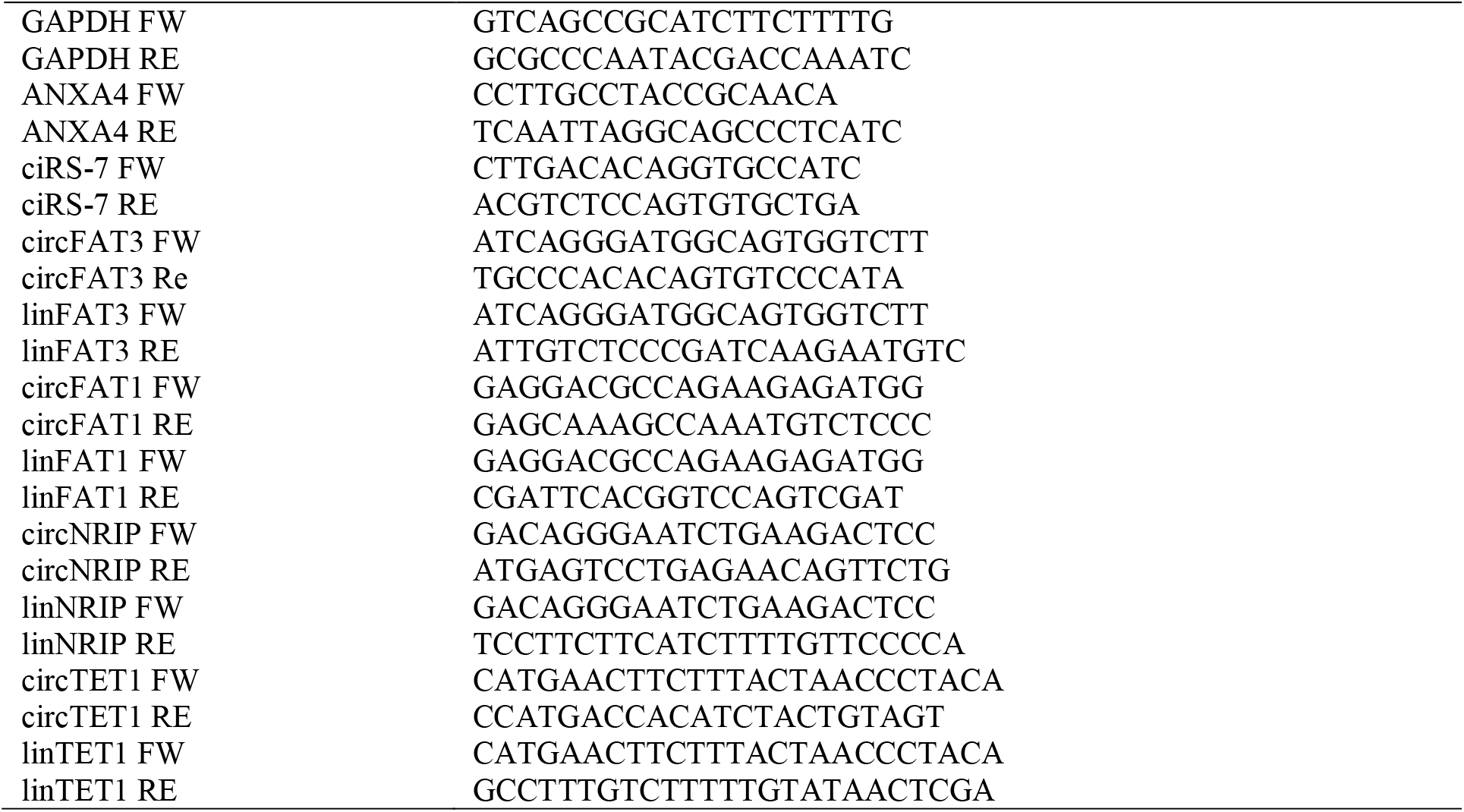
RT-qPCR primer sequences (5’>3’)

